# Directed connectivity between primary and premotor areas underlying ankle force control in young and older adults

**DOI:** 10.1101/804450

**Authors:** Meaghan Elizabeth Spedden, Mikkel Malling Beck, Mark Schram Christensen, Martin Jensen Dietz, Anke Ninija Karabanov, Svend Sparre Geertsen, Jens Bo Nielsen, Jesper Lundbye-Jensen

## Abstract

The control of ankle muscle force is an integral component of walking and postural control. Aging impairs the ability to produce force steadily and accurately, which can compromise functional capacity and quality of life. Here, we hypothesized that reduced force control in older adults would be associated with altered cortico-cortical communication within a network comprising the primary motor area (M1), the premotor cortex (PMC), parietal, and prefrontal regions. We examined electroencephalographic (EEG) responses from fifteen younger (20-26 yr) and fifteen older (65-73 yr) participants during a unilateral dorsiflexion force-tracing task. Dynamic Causal Modelling (DCM) and Parametric Empirical Bayes (PEB) were used to investigate how directed connectivity between contralateral M1, PMC, parietal, and prefrontal regions was related to age group and precision in force production. DCM and PEB analyses revealed that the strength of connections between PMC and M1 were related to ankle force precision and differed by age group. For young adults, bidirectional PMC-M1 coupling was negatively related to task performance: stronger backward M1-PMC and forward PMC-M1 coupling was associated with worse force precision. The older group exhibited deviations from this pattern. For the PMC to M1 coupling, there were no age-group differences in coupling strength; however, within the older group, stronger coupling was associated with better performance. For the M1 to PMC coupling, older adults followed the same pattern as young adults - with stronger coupling accompanied by worse performance - but coupling strength was lower than in the young group. Our results suggest that bidirectional M1-PMC communication is related to precision in ankle force production and that this relationship changes with aging. We argue that the observed age-related differences reflect compensatory mechanisms whereby older adults maintain performance in the face of declines in the sensorimotor system.

## 1. Introduction

The ability to precisely regulate ankle muscle force is a critical component of walking and postural control. Aging is commonly accompanied by an impaired ability to produce force steadily and precisely, especially during low-intensity contractions (1, 2), which can compromise balance and gait function. Ankle dorsiflexor muscles are subject to direct cortical influence (3), which may be related to the central role of the tibialis anterior in the swing phase of the gait cycle. This raises the question of whether age-related changes in dorsiflexion force control may have cortical correlates.

Prior work has demonstrated that during a range of motor tasks, older adults exhibit different brain activation patterns than those seen in younger adults (4). Motor control in older adults is typically characterized by greater and more diffuse recruitment patterns (5, 6), as well as both increases and decreases in connectivity among motor, sensory, and cognitive regions (7, 8, 9). The latter findings are particularly important in light of the notion of functional integration in task-related networks (10) as the basis for emergent behavior.

In particular, one study investigating a button press task reported that older adults exhibited greater activation of the premotor cortex (PMC) than young adults, accompanied by greater local connectivity centered around the PMC and reduced connectivity between more distant regions (9). These findings suggest that the PMC may be a key region for age-related adaptive processes during motor tasks. This notion is supported by recent findings from Michely and colleagues (8), also for button-pressing tasks, demonstrating a reverse U-shaped trajectory in PMC influences on the primary motor cortex (M1) with age. The authors suggest that this trajectory could reflect reliance on this PMC-M1 communication as an initially useful resource that eventually breaks down at later stages of aging. This study also demonstrated greater prefrontal influences on the motor system with advancing age, which is in agreement with functional MRI results showing increased prefrontal cortex (PFC) activation during motor tasks in older adults (6).

A critical discussion in the context of these findings is whether such age-related differences represent compensation - plastic adaptations that counteract structural, biochemical and/or functional decline, or de-differentiation - age differences that reflect declines in brain structure-function relationships with age (4). For the motor system, it has been argued that greater task-related activation of brain regions in older adults has a compensatory function (6, 11, 4), but little attention has been paid to the function of differences in connectivity. Although the strict classification of coupling patterns as compensation or de-differentiation requires interventional approaches - i.e., perturbation of activity to assess resulting behavioral influences - these roles can be more informally assessed by considering whether coupling patterns observed in older adults are associated with task performance and in which way. Thus, we maintain that the key feature indicating compensation vs. de-differentiation is whether the observed differences are relevant for task performance; differences that appear to be functionally relevant likely play a compensatory role, whereas non-functional differences are likely to reflect de-differentiation. One previous study evaluating such functional correlations was not able to find direct relationships between button pressing reaction times and single coupling parameters, but did show that older participants with greater net prefrontal-premotor-M1 connectivity also had faster reaction time (8) - in support of the compensation hypothesis.

Notably, this and other previous work investigating effective cortical connectivity in young and/or older adults has focused exclusively on hand and finger movements (12, 13, 7, 8), despite the broad behavioral importance of voluntary ankle muscle control. Functionally, control of the lower limbs in humans is specialized for maintaining the upright stance and walking, in contrast to reaching and grasping functions of the arm and hand. On the one hand, these differences in functional specialization for upper and lower limbs may necessitate differential cortical control mechanisms. On the other hand, there are also noteworthy commonalities between the visuo-motor demands of movements of the hand and precision control of foot position e.g. during the swing phase of walking. Here, we adopt the point of departure that the general functions of cortical regions are conserved across muscles, but that there may be differences in characteristic coupling patterns.

Dynamic causal modelling (DCM) is a framework for estimating the effective connectivity among a set of brain regions (14). DCM uses a realistic (bio-physically informed) generative model to infer plausible explanations for how electrophysiological or functional neuroimaging data were generated. Inter- and intra-regional synchronization of oscillatory activity, as documented using magnetoencephalography or EEG, is considered an important marker for integration in functional networks (15). Alpha/mu oscillations are suggested to play a role in cortical attention modulation during movement (16), whereas beta band oscillations during steady contraction may act to stabilize current motor output (17). Gamma band oscillations are also apparent in motor regions and are thought to direct changes in motor output(18, 19).

In this study, we used DCM for cross-spectral densities (CSD) to model a network comprising the PMC, M1, PFC, and posterior parietal cortex (PPC) based on EEG responses recorded from younger and older adults during a dorsiflexion force-tracing task. To quantify commonalities and differences in network coupling strengths among participants, we adopted the Parametric Empirical Bayes (PEB) framework (20, 21). Using these methods, we investigated how cortico-cortical coupling within our network was related to precision in force production; how coupling strength differed between young and older participants; and whether any associations between precision and coupling strength differed between the two age groups.

We maintained a general hypothesis that reduced force control in older adults would be associated with cortico-cortical connection strength within our network comprising M1, PMC, PPC, and PFC. We also hypothesized that that connectivity patterns in older adults would support the compensation hypothesis, i.e., that differences in coupling patterns observed in older adults would be relevant for (correlated with) task performance.

## 2. Methods

### 2.1. Participants

Fifteen younger (mean 22.1 ± 1.7, range 20 to 26 yr, 8 female) and fifteen older adults (mean 68.3 ± 2.7, range 65 to 73 yr, 8 female) with no reported neurological or neuromuscular disorders and Mini Mental State Exam (MMSE) scores ≥ 26 were recruited for this study by convenience sampling. Twenty-one participants were right leg dominant, 2 were left-leg dominant, and 7 were equally right- and left leg dominant (Waterloo Footedness Questionnaire; Elias and Bryden, 1998) (22).

All participants provided written, informed consent prior to experiments, and the study was approved by the ethics committee for the Capital Region of Denmark (approval number H-16021214). Experiments were conducted in accordance with the Declaration of Helsinki. Some of the data from this study have been previously published in a paper investigating cortico-muscular coherence between the scalp EEG and electromyographic recordings (23). Participants also performed other motor tasks during the same experiments which are reported elsewhere (24).

### 2.2. Force tracing task

Participants performed a simple force tracing task, challenging precision in ankle muscle torque production (Figure 1B) (23).While participants were seated in a chair with their left leg fastened to a force pedal with a built-in strain gauge, the leg was positioned to 120° dorsiflexion at the ankle, 115° flexion at the knee, and 90° flexion at the hip. Participants initially performed three attempts at a maximal voluntary dorsiflexion contraction (MVC). The peak torque attained across these attempts was used to calculate 10 % MVC as the target level for the tracing task.

**Figure 1:**
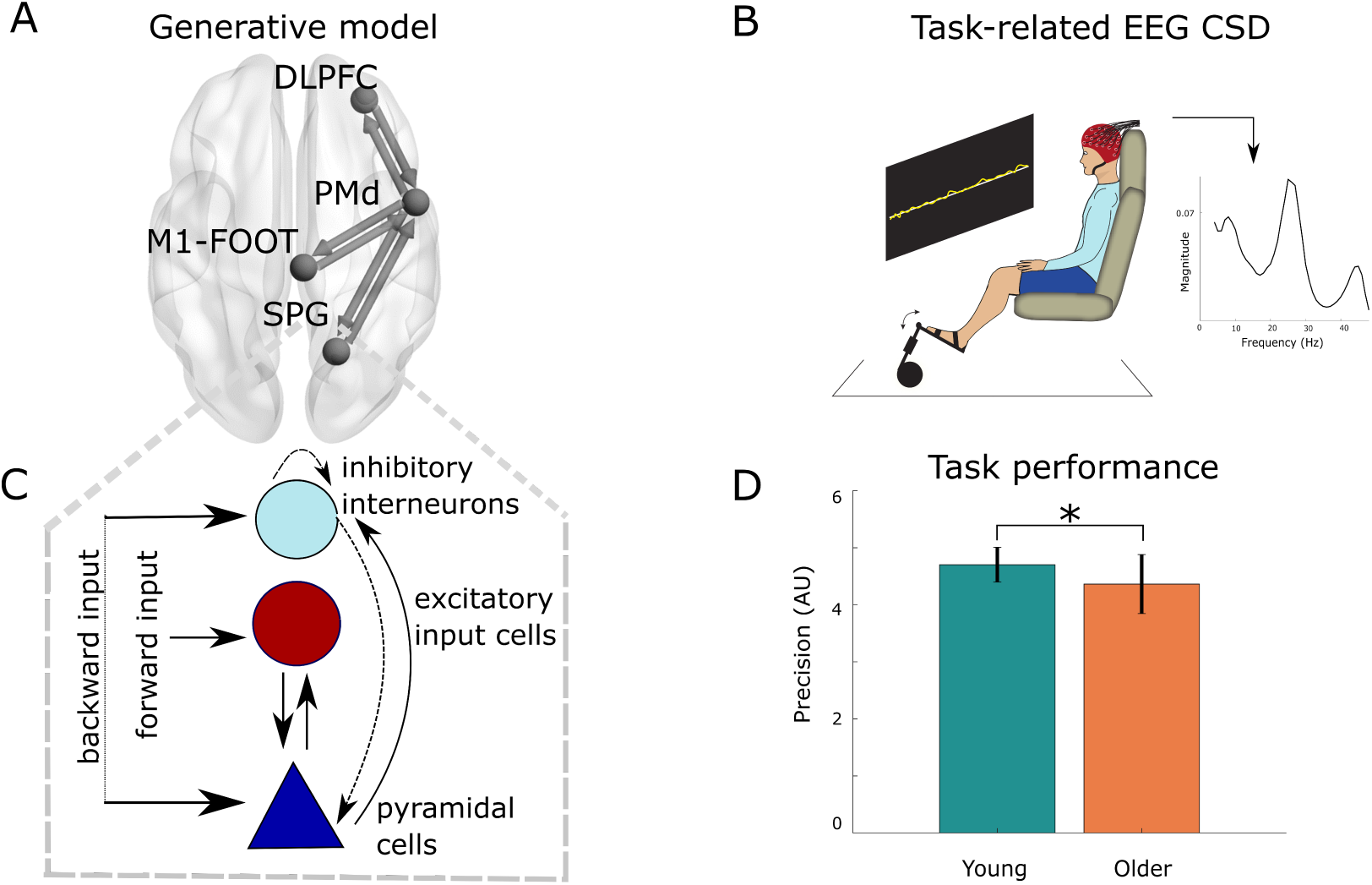
Methods overview. Effective connectivity in a uni-hemispheric network for the contralateral hemisphere comprising the dorsolateral frontal cortex (DLPFC), the foot area of the primary motor cortex (M1-FOOT) and dorsal premotor cortex (PMd), and the superior parietal gyrus (SPG) (A) was inferred using dynamic causal modelling of EEG auto- and cross-spectral densities (CSDs) during a force tracing task using the left ankle muscles (B). Activity in each source was described using the local field potential (LFP) neural mass model (C) and passed through lead fields to generate auto- and cross-spectra for principal EEG modes. We investigated whether network parameters differed in older and younger adults and were related to precision during the force tracing task (D). Dotted lines in C indicate inhibitory connections. * indicates p *<* 0.05. AU, arbitrary units.

The goal of the force tracing task was to trace the target level as precisely as possible for two minutes. Participants were aided by a projection of the target level (shown as a horizontal line) as well as their actual torque production (shown as a real-time trace) onto the wall in front of them (screen size 125 cm by 167 cm). Task performance was assessed based on the root mean square (RMS) error, i.e., the deviation of actual torque from the target level over the two minutes. RMS values were first log transformed to normalize their distribution, then multiplied by −1 to render their interpretation more straightforward, such that higher values corresponded to better performance (greater precision). Age-group differences in log-transformed task performance were tested using an unpaired t-test.

### 2.3. EEG

#### 2.3.1. Recording

Participants were fitted with an EEG cap accommodating 64 active electrodes (BioSemi, Amsterdam, The Netherlands) positioned according to the 10-20 system. Signals were recorded using ActiView software (v6.05) with a sampling frequency of 2048 Hz. Common mode sense and driven right leg electrodes served as the online reference as per BioSemi design. Before recordings, participants were reminded to relax face and neck muscles to minimize signal artifacts. We also monitored EEG signals during recordings to ensure that electrode offset was maintained below 25 *µ*V. Signals were recorded continuously for ∼ 2 minutes.

#### 2.3.2. Pre-processing

EEG data was pre-processed using EEGlab software (v14.0.0) in MATLAB (vR2016b). EEG signals were visually inspected with the aim of identifying and removing noisy channels before signals were re-referenced to an average reference. Signals were then band-pass filtered from 0.5 to 48 Hz and down-sampled to 256 Hz. Periods of marked, high-amplitude deviations (e.g. from strong muscle artifacts or periodically loose electrodes) were also removed. Independent components analysis decomposition was then performed using the runica algorithm, and components displaying topological and spectral qualities indicating eye blinks and saccades were removed (25). Finally, removed channels were interpolated using the standard spherical splines approach. Further analysis was performed in SPM12 (r7487). EEG signals were segmented into 1-s epochs, resulting in 116-125 epochs per participant, and channel coordinates were transformed into corresponding coordinates in MNI space.

#### 2.3.3. Dynamic Causal Modeling

Dynamic causal modelling (DCM) is a method for inferring the effective, task-related connectivity between a set of brain regions from electrophysiological responses (26). Here, we used DCM for CSD, designed to describe steady-state network dynamics (27, 28). DCM for CSD explains complex cross-spectra as generated by a network of dynamically coupled sources, where each source is represented by a neural mass model (29). The foundation of this DCM is a set of differential equations that characterize the response of populations of neurons in each source to endogenous and exogenous input. These equations are augmented with a spatial forward model to describe how responses of neuronal populations are translated into EEG recordings. This generative model thus attempts to describe the dynamics of hidden states - interactions between neuronal populations - and how these states effectuate the observed CSDs. Bayesian estimation (inversion) of the ensuing DCM (30) provides the posterior distribution of model parameters that are obtained through maximizing the model’s negative variational free energy, i.e. accuracy minus complexity (penalizing deviations from priors). Our primary parameters of interest in this study are extrinsic connection strengths, i.e., connections between network regions.

#### 2.3.4. Neural mass model

In DCM for EEG, source activity can be represented by different neurobiologically informed neuronal population models (29). We adopted the convolution-based local field potential (LFP) neural mass model, which describes source activity as the result of interactions between populations of inhibitory interneurons, excitatory input cells, and excitatory pyramidal cells, inspired by the laminar structure of the cortex (31). Our choice of the LFP neural mass model was motivated by the minimal model approach, i.e., we chose the simplest neural mass model allowing access to the parameters of interest, namely connection strengths between sources (29). Figure 1C illustrates the LFP model. Each hidden state parameterizes the firing rate of a neuronal population and depends on the average pre-synaptic inputs, post-synaptic membrane potential, and constants describing biophysical membrane properties. Forward connections between network regions can be interpreted as implementing a strong driving effect, whereas backward connections have more modulatory effects on their target populations. For a full mathematical description, we refer to (29).

#### 2.3.5. Network regions

In the DCM for EEG framework, network architecture is determined a priori. Our full hypothesized network comprised a uni-hemispheric network for the contralateral hemisphere containing the foot area of M1 and the dorsal PMC (PMd), the superior parietal gyrus (SPG), and dorsolateral frontal cortex (DLPFC). The choice of these regions was carefully motivated based on previous fMRI studies of ankle movements (32, 33, 6, 34). Across different tasks involving visually cued or guided (32, 33, 34) and challenging cyclical (6) ankle movements, these studies showed that a core motor network involving M1, PMC and supplementary motor area (SMA) was activated. For our network, we chose to include M1 and PMC, but omitted SMA on the grounds that it is thought to play a more central role in self-generated movement, in contrast to the more prominent role of PMC in externally-guided movement (35). Also, the proximity of SMA to the foot area of M1 would likely make it difficult to distinguish between the two areas reliably. Some of these studies also demonstrated activation in prefrontal areas (32, 33, 6), and results from one of these studies also indicated greater, compensatory PFC activation in older adults (6). Thus, we also included influences from the DLPFC on the core motor circuit in our network. Finally, we wanted to include a posterior parietal region to account for sensorimotor interactions related to the use of visual and proprioceptive information to guide motor behavior. Prior work on cyclical and visually guided ankle movements has indeed demonstrated PPC activation (36, 6, 34), so we also included the SPG region in our network. The full DCM model thus included forward connections from SPG to PMd; DLPFC to PMd; and PMd to M1; and backward connections from PMd to SPG; PMd to DLPFC; and M1 to PMd (37) (Figure 1A). Altogether, we regarded this network as the optimal compromise between selecting the simplest (limiting the number of model parameters) and ostensibly most relevant network for the task. We used MNI coordinates for our regions of interest reported in previous fMRI studies (33, 6) (Table 1). Brain network figures were generated using BrainNetViewer (38).

**Table 1:**
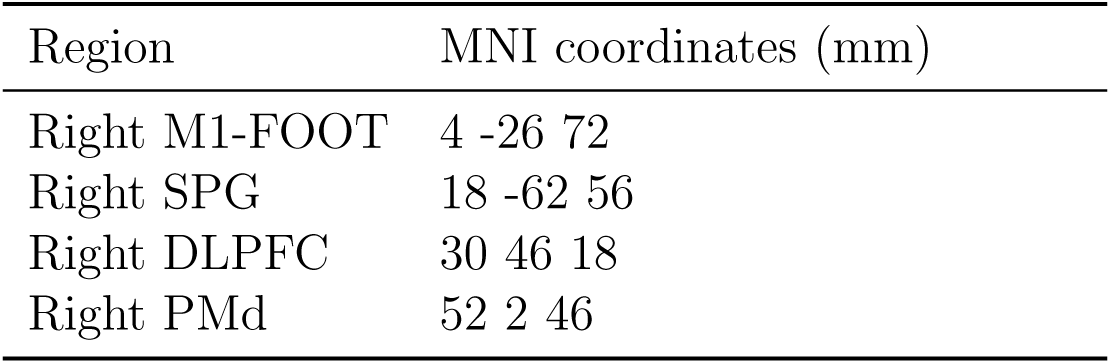
Prior coordinates for source locations. M1-FOOT, primary motor cortex, foot area; SPG, superior parietal gyrus; DLPFC, dorsolateral frontal cortex; PMd; dorsal premotor cortex. Coordinates are from (33, 6).

#### 2.3.6. Spatial forward model

Inverting a DCM for CSD subsumes inverting a classical spatial forward model to map how source activity is translated into predicted EEG data. We used the default SPM forward model, which comprises a linear translation from neuronal states to EEG signals. Our source space was modelled using SPM’s template head based on the MNI brain. When using this template, EEG electrode positions are transformed to match the template head (39), so even in the case of discernable differences between participant and template heads, the result should be reasonable. The forward model was constructed using the boundary element method (BEM). A Bayesian approach using the multiple sparse priors (MSP) algorithm was applied to invert the forward models (40), projecting sensor data to source space.

#### 2.3.7. Spectral estimates

The features to be predicted by this DCM were auto- and cross-spectra between the 8 first principal (eigen)modes of EEG channel mixtures. These principal modes were used to reduce the dimensionality of the data. Cross-spectral densities were computed using Bayesian multivariate autoregressive modelling (41) with the default model order of 8 (28). We chose a broad frequency range from 4-48 Hz to account for alpha, beta, and low gamma activity (42).

After encountering local minima during model inversion, we adjusted the hyperprior for the expected precision of the data (hE=18) to increase reliance on achieving accurate fits at the expense of prior values (43). Also, due to the presence of spectra with rather prominent features (not conforming to the 1/f assumed by DCM), we set the prior of the neural innovation to assume a flat spectrum. For all other priors, we used default settings (28).

#### 2.3.8. Second level analysis using PEB

After estimating all connectivity parameters of interest (full DCMs) for each participant, we used the PEB framework (44, 21) to characterize commonalities between subjects in network connection strengths, as well as differences due to age-group; task performance; and interaction between age-group and performance. A notable advantage of this framework, as opposed to classical ANOVA, is that it takes not only the mean, but also the uncertainty of individual connection strengths into account. This entails that participants with more uncertain parameter estimates will be down-weighted, while participants with more precise estimates receive greater influence (21).

The PEB approach involves (i) estimating group level parameters using a general linear model (GLM) that divides inter-subject variability into regressor effects and unexplained random effects, followed by (ii) comparison of different combinations of these parameters to identify those that best explain commonalities and differences in connectivity due to age group and performance (Bayesian model comparison).

To do this, we used an exhaustive comparison of all possible GLMs where one or more second-level parameters were turned off (fixed at prior of zero) to prune away parameters that did not contribute to the model evidence (Bayesian model reduction). Model evidence is also quantified at the second level as negative variational free energy, which here is the sum of DCM accuracies for all participants minus complexity due to fitting both the DCMs and the GLM. The automatic search was performed under the assumption that all reduced models were equally probable and that the full DCM only contained physiologically plausible parameters (21). Finally, we calculated a Bayesian Model Average (BMA) over the GLMs from the final iteration of the search, in which parameters were weighted by the models’ posterior probabilities.

We used SPM’s default prior values for inter-subject variability and second level parameters and took only the extrinsic connectivity values (i.e., forward and backward connections between network regions) to the second level, as we were only interested in the effect of regressors (i.e., age group; task performance and interactions between age group and task performance) on these parameters. Thus, we report as our main outcomes: mean connection strengths, their uncertainties, and probabilities across all participants; differences in connectivity due to age-group; relationships between connectivity and task performance; and relationships between connectivity and task performance that differ by age-group (interactions). These interpretations were enabled by mean-centering regressors before entering them into the GLM. As the statistics performed here are Bayesian, we chose to threshold averaged model parameters to include those with posterior probabilities *>* 95 % (i.e. strong evidence) when comparing models with and without the parameter based on their free energy. In scatter plots illustrating associations between task performance and connectivity strength, we used updated DCM values where (log-scaling) parameters had been re-evaluated using the group average connection strengths as priors to obtain the most robust estimates (21).

## 3. Results

### 3.1. Task performance

A comparison of task performance between the two age-groups revealed that precision was lower for the older than for the younger group (*t*_28_=-2.193, p=0.037, Figure 1D), indicating less accuracy in dorsiflexion force production in older participants (as previously reported for these participants in (23)).

### 3.2. EEG spectra and DCM fits

The principal modes in our EEG data showed auto- and cross spectra often containing one or more marked peaks in alpha and beta bands. Figure 2 shows exemplar spectra and predictions from the DCM inversion from a single participants data. As illustrated, the DCMs were able to capture the most prominent spectral elements. However, more intricate spectra with multiple features proved more challenging to fit, given that secondary peaks were sometimes neglected by the models (e.g. mode 4 to 2 in Figure 2). Full DCMs were successfully fitted for all 30 participants without indications of early convergence. Mean variance explained by the fitted DCMs was 93.5 % (range 76.8 - 98.1 %).

**Figure 2:**
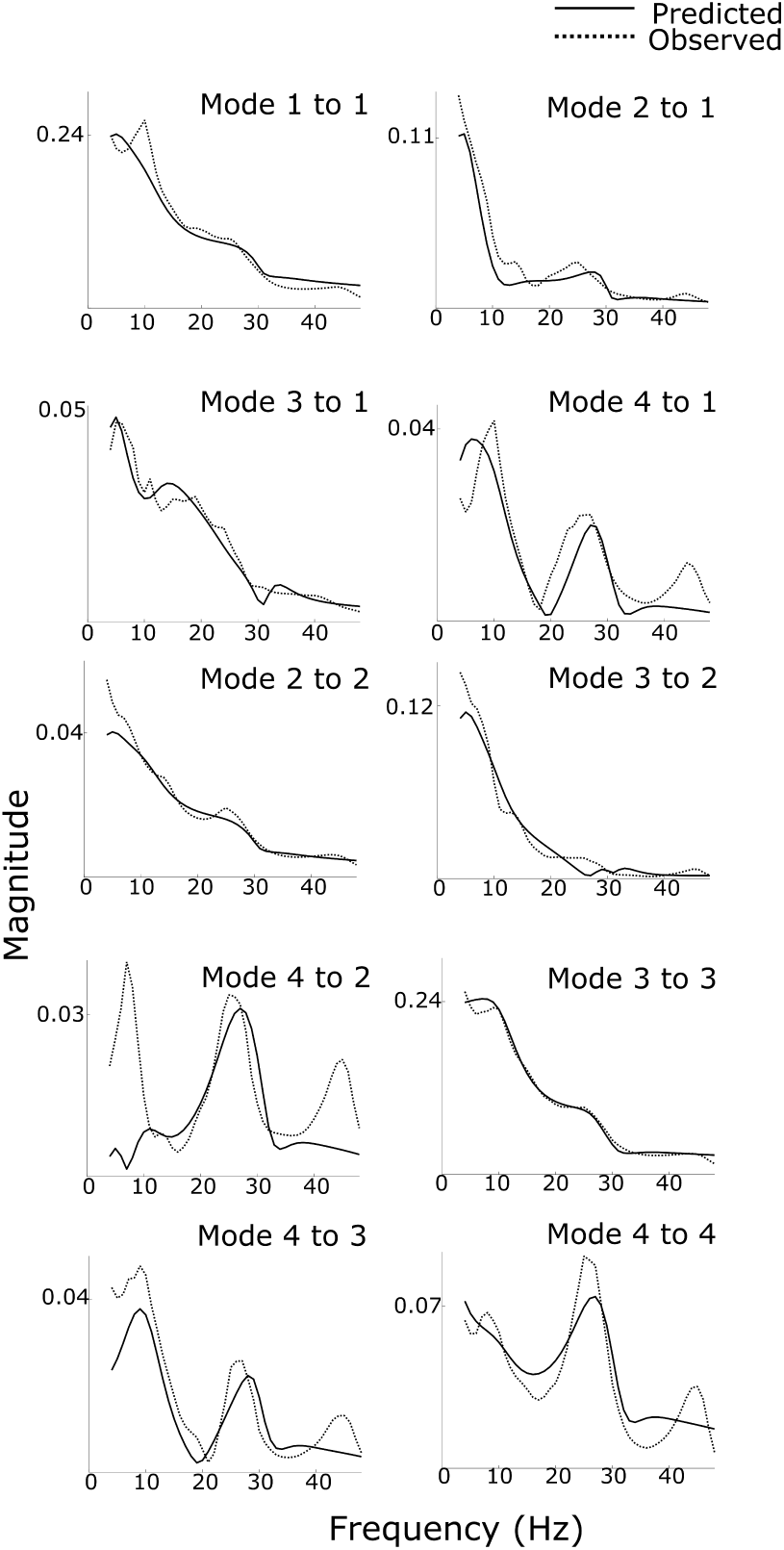
Model predictions. Dynamic causal model (DCM)-predicted and observed auto- and cross spectra for the first 4 principal EEG modes for a single exemplar participant.

### 3.3. PEB

Using PEB, we estimated extrinsic connectivity strengths and their uncertainties at the second (group) level to determine commonalities between all participants; effects of age-group; effects of performance; and age-group performance interactions. As we did not have strong hypotheses regarding precisely where in our network these effects would be expressed, we used BMR to prune away connections that did not contribute to the model evidence, followed by BMA to compute a weighted average of the parameters in the final BMR iteration. Figure 3 shows this BMA of group-level estimates of network connection strengths, their uncertainties, and posterior probabilities thresholded for *>* 95% posterior probability.

**Figure 3:**
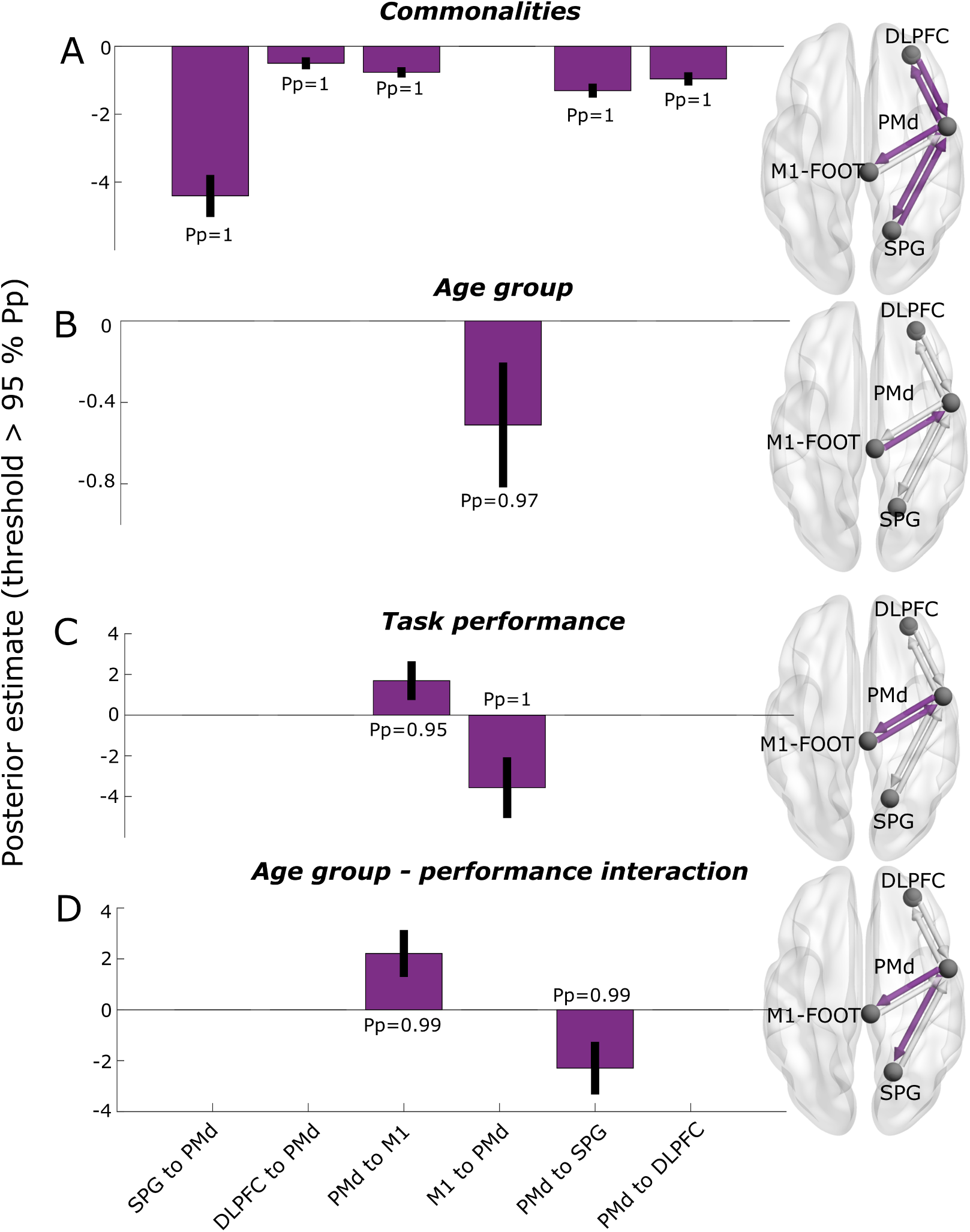
Posterior parameter estimates. Bar plots are thresholded for parameters *>* 95 % posterior probability (Pp). Purple arrows in brain figures indicate connection parameters exhibiting Pp *>* 95 %. SPG, superior parietal gyrus; DLPFC, dorsolateral frontal cortex; PMd, dorsal premotor cortex; M1/M1-FOOT, foot area of primary motor cortex.

#### 3.3.1. Commonalities

The BMA of parameters relating to commonalities across all participants (Figure 3A) indicated that 5 out of the 6 DCM connections were conserved across subjects, exhibiting a probable, non-zero group mean. However, the backward connection from M1 to PMd showed a group mean of zero when considering all participants together. The commonality parameter estimates indicate mean connection strengths, and the negative signs here are due to the procedure of parameterizing connectivity values in terms of log-scaling parameters. Negative values indicate that the estimated connectivity is down-scaled relative to the prior mean.

#### 3.3.2. Age-group

Figure 3B shows a probable effect of age-group on the backward connection from M1 to PMd. The negative value of this parameter indicates that the strength of this connection was weaker in older than in younger participants. When we plotted individual connectivity values (unitless log-scaling parameters) for each age-group (see Figure 4A), it became clear that the mean value of zero for this parameter across all participants was the result of connectivity values for each age group roughly positioned on each side of zero. No other effects of age-group on connectivity were present thresholded at *>* 95% posterior probability.

**Figure 4:**
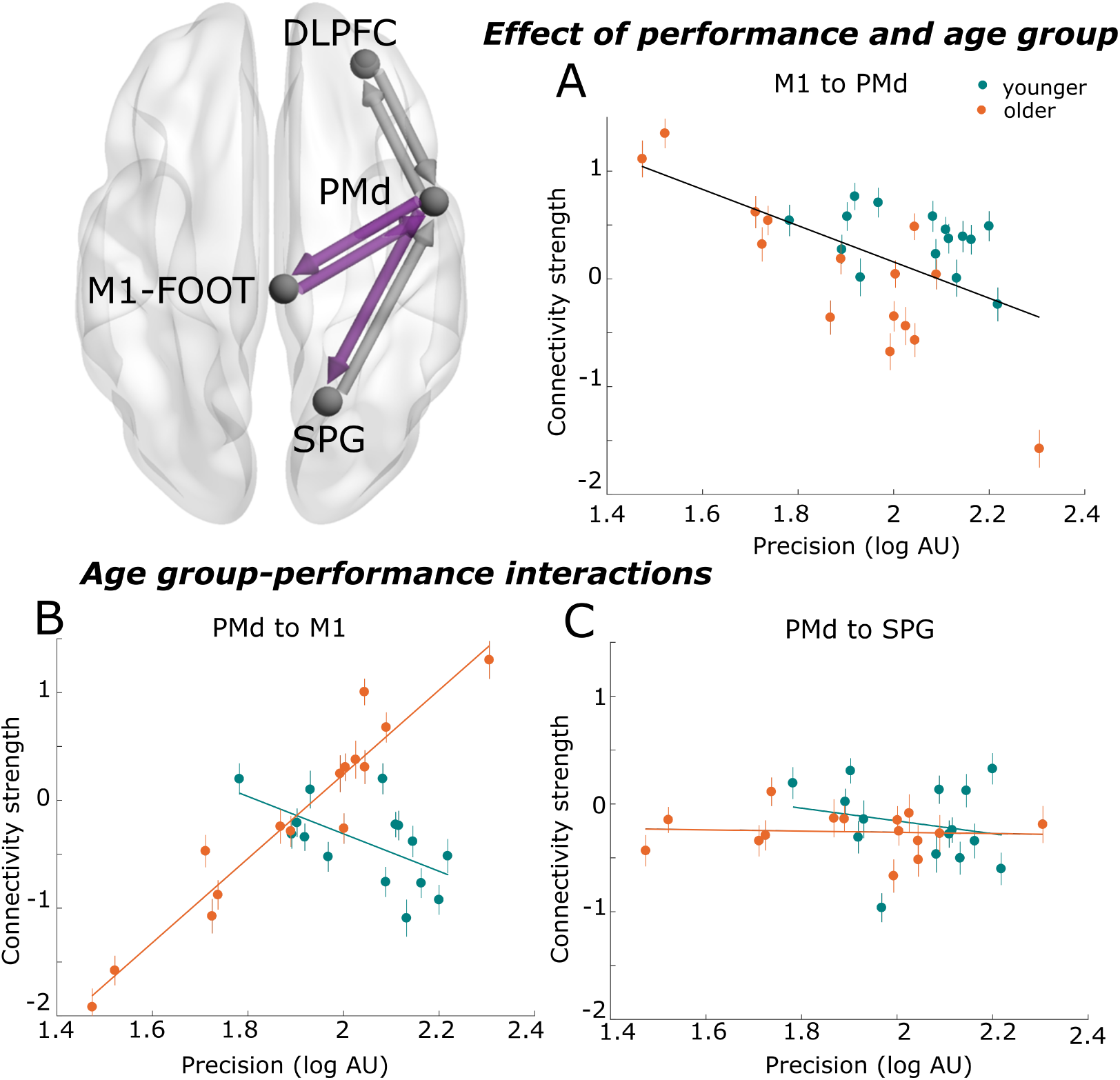
Main effects and interactions. Relationship between precision in dorsiflexion force production and M1 to PMd connectivity (A); PMd to M1 connectivity (B); and PMd to SPG connectivity (C) for the two age groups. Connectivity strengths are shown as unitless log-scaling parameters. Error bars indicate parameter estimate uncertainties. SPG, superior parietal gyrus; DLPFC, dorsolateral frontal cortex; PMd, dorsal premotor cortex; M1/M1-FOOT, foot area of primary motor cortex; AU, arbitrary units.

#### 3.3.3. Task performance

We also detected an effect of performance (precision) on the backward connection from M1 to PMd. The negative sign of this estimate reflects that stronger M1-PMd connectivity was associated with worse performance across all participants. Figure 4A shows M1 to PMd connectivity values plotted as a function of performance and by age group, illustrating two interesting components of this relationship: a clear association between lower coupling strength and better performance; and lower coupling strength in older than younger participants. In addition, a non-trivial effect of performance on the forward connection from PMd to M1 was present (Figure 3C), but this effect must be interpreted in light of the interaction detected for this parameter.

#### 3.3.4. Age-group performance interactions

The forward PMd to M1 connection also showed a probable effect of age-group performance interaction (Figure 3D). Figure 4B clarifies this interaction as different relationships between performance and connectivity strength for the two age groups; for the young group, stronger connectivity was associated with worse performance, whereas for the older group, stronger connectivity was associated with better performance. The strength of the association present in the older group appears quite remarkable and was apparently driving the effect of performance on this parameter (Figure 3C).

Similarly, the backward PMd to SPG connection showed an interaction effect (Figure 3D), also suggesting distinct associations between performance and connection strength in older and younger participants for this connection (Figure 4C). The slope of this association for young participants was negative, indicating that participants with stronger PMd to SPG connectivity also exhibited lower precision. For the older group, however, the slope of this relationship was largely flat, suggesting a lack of association.

## 4. Discussion

Our results showed that the strength of directed interactions between PMd and M1 were related to precision in ankle force production and differed in older and younger adults. In the young group, stronger bidirectional M1-PMd coupling was associated with worse precision on the force tracing task. In the older group, the backward M1 to PMd connection followed the same pattern (i.e. stronger coupling associated with worse precision), but coupling was weaker than in the young group. Coupling strength for the forward PMd to M1 connection did not differ between age groups, but showed the reverse relationship with performance for older participants, i.e., stronger PMd-M1 coupling was associated with better performance. We suggest that these age-related differences - in light of their ostensible functionality - reflect compensatory mechanisms whereby older adults counteract sensorimotor declines to maintain performance.

### 4.1. Task performance

As presented in our previous paper (23), we observed lower precision in dorsiflexion force control in older than in younger participants. This is in line with prior work demonstrating age-related decrements in the ability to produce and regulate force steadily in lower leg muscles (1, 45). Interestingly, we found that the degree of precision in ankle force production was dependent on several cortical connectivity parameters in older and young participants: PMd-M1 interactions, and feedback influences from PMd to SPG.

### 4.2. Stronger bidirectional M1-PMd coupling was associated with worse precision in young adults

Based on studies of reaching and grasping movements, the PMC is thought to play a central role in planning and selecting motor responses based on external cues (46, 47). Premotor communication with the PPC facilitates the use of visual and proprioceptive information to formulate motor commands, which are executed through coupling with M1 (48). The PMC also has close connections with the PFC (49), which mediate cognitive influences on the selection and modulation of motor responses. Thus, forward (driving) PMd to M1 connections likely serve to direct movements based on executive influences and sensory information, whereas backward (modulatory) M1 to PMd connections likely provide contextual guidance aiding ongoing adjustments of motor plans.

We found that the strength of coupling from PMd to M1 and from M1 to PMd was negatively related to force precision for young participants, i.e., those with weaker coupling performed better. Based on previous work on the role of these regions in reaching and grasping movements, this was surprising. Given the putative role of the PMd in using external cues to guide motor output, we might expect that stronger forward PMd to M1 coupling would promote better performance. Likewise, stronger feedback from M1 to PMd might be expected to improve performance, as updating PMd with information about M1 output could facilitate adjustments based on changes in the current state. However, when considering the nature of the force-tracing task, stronger M1-PMd coupling may not be advantageous in this context. In this type of steady state task with stable properties, the need for PMd monitoring of motor output state may be minimal and may even represent a suboptimal strategy leading to redundant adjustments in motor output.

In support of this, we also observed a negative association between performance and PMd-SPG coupling strength in the young group. Whereas forward communication from SPG to PMd likely codes a spatial reference frame for the intended movement (at least for reaching and grasping) (50), the corresponding backward communication may serve to update sensory integration with contextual information about developing plans. This negative association may thus also reflect the type of task we used where online modulations of movement goals (requiring continuous SPG updating) are unfavorable for precise performance.

Taken together, these negative associations likely reflect that good young performers make use of alternative routes of communication for successful force control. One possible explanation - despite our hypothesis that the force-tracing task would be heavily reliant on guidance from external cues - is that the sensory monitoring required for this task was limited, and that younger participants performed the task to a greater extent based on internal representations. Specifically, it could be the case that young participants were more reliant on communication within the so-called intrinsic circuit comprising the SMA, basal ganglia and M1, which has been suggested to control self-generated movements, rather than the extrinsic circuit containing PMC, cerebellum, M1, and parietal areas, which is thought to control movement guided by external cues (51, 52). Thus, when tracing the force target, participants may have quickly learned to predict the result of ankle movements on the torque signal such that movements were controlled by an internal model - preferentially through the intrinsic circuit - without a considerable dependency on visual and somatosensory feedback. In any case, our results suggest that young participants exhibiting better precision were not reliant on the cortical coupling parameters included in our network for successful force control.

Another important consideration is whether these negative associations between coupling strength and performance in the young group could reflect differences in the control of lower and upper extremities. Data from our study cannot be used to specifically compare coupling patterns during hand- and leg-based tasks, so we can only speculate. Nevertheless, the functional specialization of hand control for reaching and grasping is in some aspects distinct from the specialization of lower limbs for standing and walking, which may entail that leg movements are coordinated by different patterns of effective connectivity - e.g. that the PMC is less involved in visually guided ankle than hand movements.

On the other hand, there are also important similarities between foot and hand control: as is the case for arm and hand movements, foot position during walking must be fine-tuned based on visual and proprioceptive feedback to avoid obstacles and accommodate changes in terrain. For example, during the swing phase of walking, the toes typically clear the ground very precisely with a variability of less than 4 mm (53), which may be considered a control challenge on par with the precision grip of the hand. In addition, the notion of an analogous role for the PMC in the control of lower extremities is supported by gait studies demonstrating PMC activation during imagined and real walking tasks, including paradigms emphasizing precision (54, 55, 56). Thus, the negative correlations between PMd-M1 connectivity and performance may primarily reflect task-specific strategies rather than differences in control of upper- and lower extremities. Future work comparing coupling parameters during the same task performed with hand and ankle muscles is necessary to clarify these issues.

### 4.3. Lower feedback coupling from M1 to PMC in older adults

For the feedback connection from M1 to PMd, older participants followed the pattern observed in younger participants, i.e. stronger coupling was associated with lower force precision. However, the older group exhibited lower connectivity strength compared to the young group for this parameter. A possible interpretation of this difference is that the weaker coupling observed in older adults represents a compensatory mechanism, in light of the finding that weaker coupling was associated with better performance across groups. If we characterize compensation based on the functional relevance of the observed deviations, then this finding could reflect an adaptive response to maintain performance in the face of structural, functional and/or biochemical declines in the sensorimotor system (57, 58, 4, 59). The ostensible weakening of coupling with advancing age may thus be a useful strategy to improve performance. However, precision in the older group was still lower than the younger group, suggesting that this strategy was not able to compensate fully.

### 4.4. Stronger forward PMC-M1 coupling associated with better performance in older adults

Although the strength of the forward PMd-M1 coupling did not differ between age groups, older participants exhibited a strong, positive association between coupling strength and precision, in contrast to the reverse, negative association observed in young participants. This discrepancy suggests age-related differences in the reliance on PMd-M1 communication for task performance. Better performers in the older group appear to have capitalized on PMd-M1 influences to achieve good performance, which, in conjunction with a corresponding attenuation of feedback from M1 to PMd, may be a reflection of greater dependency on feedforward communication between these regions. This might be understood in terms of the predictive coding framework; it has for example been suggested that the aging brain becomes progressively optimized to generate increasingly accurate predictions of the environment (60). These authors discuss that the greater sensory experience that older adults possess may contribute to the refinement of generative models, more efficient predictive control and thus reduced reliance on feedback communication. Note however that we have only observed this pattern for PMd-M1 communication, and not in other regions in our network. Another possible interpretation is that an increased reliance on influences from PMd on M1 could reflect greater attention directed towards task performance in good older performers (61). We argue in any case that this coupling pattern may also represent a compensatory mechanism, at least for the good older performers able to make use of this connectivity to support performance. Taken together, our results suggest that PMd-M1 communication becomes more relevant for precision control of the ankle joint with aging.

### 4.5. The PMC and PFC as compensatory resources

Of interest, the most robust effects of age group and performance were centered around M1-PMd interactions. The PMC has been previously highlighted as a key region for adaptive processes in the motor system, particularly in the case of reduced M1 function. Behaviorally meaningful adaptations have been demonstrated in PMC after M1 lesions in animals and humans (62) and non-invasive brain stimulation suppressing M1 excitability (63), indicating that functional compensation may depend on PMC activity. Further, other studies have demonstrated age-related increases in PMC activation during motor tasks (5, 9, 64), and some of this work has additionally suggested that this activation may become increasingly useful with aging (64). Our results expand upon this previous work by demonstrating that both forward driving and backward modulatory PMd-M1 connections exhibit distinct age-related differences that are directly linked to behavior.

Also of interest, we did not detect any probable effects of age-group or performance on coupling between the prefrontal and premotor regions. This is somewhat inconsistent with prior studies indicating that the PFC may be an important compensatory resource for motor control in older adults (5, 8, 4). The most obvious explanation for this discrepancy is the difficulty of the task utilized. Previous work suggests that the reliance on PFC influences in older adults may first emerge with increasing task difficulty (6), which could mean that for the relatively simple force tracing task we used, adjustments in coupling between core motor areas were sufficient for older adults to perform the task.

### 4.6. The role of descending drive in precision force control

It has been suggested that force fluctuations are a consequence of low frequency (delta band) oscillations embedded in neural drive to the active muscles, and that an age-related increase in this oscillatory input leads to lower force control in older adults (1, 65). Given that the regulation of these oscillations appears to be supra-spinal in origin (66), the age differences we observed in PMd-M1 coupling may play a role in modulating this oscillatory input to motor neuron pools with aging. As both PMd and M1 have corticospinal projections to the spinal cord (46), interactions between these regions are in any case anatomically suited to perturb corticospinal activity and thus affect descending drive to the spinal motor neuron pools.

## 5. Conclusion

Our results showed that directed cortical interactions between PMd and M1 were associated with ankle force precision and differed by age group. For the young group, bidirectional PMd-M1 connection strength was negatively related to task performance: stronger backward M1-PMd and forward PMd-M1 coupling was associated with worse precision. Notably, the older group showed departures from this pattern. For the PMd to M1 coupling, there were no age group differences in coupling strength, but within the older group, stronger coupling was associated with greater precision. For the M1 to PMd coupling, older adults followed the same pattern as young adults - with stronger coupling accompanied by worse performance - but coupling strength was lower than in the young group. We argue that the observed age-related differences in coupling patterns reflect useful adaptive mechanisms through which older adults maintain performance in the face of declines in the sensorimotor system. These results add to the discussion of the cortical control of ankle muscles and adaptive plasticity with aging.

## Acknowledgements

We would like to thank the participants for their engagement in this study. We are also very appreciative of DCM assistance from Rosalyn Moran and the SPM team.

